# Genetic evidence for two carbon fixation pathways in symbiotic and free-living bacteria: The Calvin-Benson-Bassham cycle and the reverse tricarboxylic acid cycle

**DOI:** 10.1101/363564

**Authors:** Maxim Rubin-Blum, Nicole Dubilier, Manuel Kleiner

## Abstract

Very few bacteria are able to fix carbon via both the reverse tricarboxylic acid (rTCA) and the Calvin-Benson-Bassham (CBB) cycles, such as symbiotic, sulfur-oxidizing bacteria that are the sole carbon source for the marine tubeworm *Riftia pachyptila*, the fastest growing invertebrate. To date, this co-existence of two carbon fixation pathways had not been found in a cultured bacterium and could thus not be studied in detail. Moreover, it was not clear if these two pathways were encoded in the same symbiont individual, or if two symbiont populations, each with one of the pathways, co-existed within tubeworms. With comparative genomics, we show that *Thioflavicoccus mobilis*, a cultured, free-living gammaproteobacterial sulfur oxidizer, possesses the genes for both carbon fixation pathways. Here, we also show that both the CBB and rTCA pathways are likely encoded in the genome of the sulfur-oxidizing symbiont of the tubeworm *Escarpia laminata* from deep-sea asphalt volcanoes in the Gulf of Mexico. Finally, we provide genomic and transcriptomic data suggesting a potential electron flow towards the rTCA cycle carboxylase 2-oxoglutarate:ferredoxin oxidoreductase, via a rare variant of NADH dehydrogenase/heterodisulfide reductase. This electron bifurcating complex, together with NAD(P)+ transhydrogenase and Na+ translocating Rnf membrane complexes may improve the efficiency of the rTCA cycle in both the symbiotic and the free-living sulfur oxidizer.

**Importance:** Primary production on Earth is dependent on autotrophic carbon fixation, which leads to the incorporation of carbon dioxide into biomass. Multiple metabolic pathways have been described for autotrophic carbon fixation, but most autotrophic organisms were assumed to have the genes for only one of these pathways. Our finding of a cultivable bacterium with two carbon fixation pathways in its genome opens the possibility to study the potential benefits of having two pathways and the interplay between these pathways. Additionally, this will allow the investigation of the unusual, and potentially very efficient mechanism of electron flow that could drive the rTCA cycle in these autotrophs. Such studies will deepen our understanding of carbon fixation pathways and could provide new avenues for optimizing carbon fixation in biotechnological applications.

## Observation

Primary production by autotrophic organisms drives the global carbon cycle. Currently, seven naturally occurring pathways for inorganic carbon fixation are known in autotrophic organisms (1). The dominant carbon fixation pathway used by plants, algae, and many bacteria is the Calvin-Benson-Bassham (CBB) cycle. The six more efficient, alternative pathways are limited to autotrophic microbes that live in reducing habitats, due to the oxygen sensitivity of these alternative pathways (2, 3). Only a handful of autotrophic organisms have more than one carbon fixation pathway: The sulfur-oxidizing symbionts of marine tubeworms such as *Riftia, Escarpia, Tevnia and Lamellibrachia* (the symbionts of these hosts are closely related to each other) have and express both the oxygen-tolerant CBB cycle and the oxygen-sensitive reverse tricarboxylic acid (rTCA) cycle (4–8). Only a few free-living bacteria may have the genes for both cycles, such as the large sulfur bacteria, *Beggiatoa* and *Thiomargarita* spp., in which all CBB cycle genes and some rTCA cycle genes were found to co-exist in their genomes (9–11). The CBB cycle in the symbionts and the large sulfur bacteria is potentially more energy efficient than the classical version of the CBB cycle based on the replacement of the fructose-1,6-bisphosphatase with a pyrophosphate dependent enzyme (9, 10, 12, 13). In addition, it is likely that the interplay between the CBB and rTCA cycle under fluctuating redox conditions contributes to the high efficiency of carbon fixation in tubeworm symbioses (4, 5, 14), and consequently to the extremely high growth rates of tubeworms, which grow faster than any other known invertebrate (15).

Given that tubeworm symbionts and large sulfur bacteria could not yet be cultivated, it was not possible to investigate the co-occurrence of two carbon fixation cycles in detail to better understand the biochemical and physiological mechanisms that enable the interplay between these two pathways. In this study, we sequenced the genome and transcriptome of the symbiont from the tubeworm *Escarpia laminata* and compared its genome to those of other tubeworm symbionts and free-living microbes. These comparisons led us to discover the presence of cooccurring CBB and rTCA cycles in the genome of a cultured bacterium.

### Co-occurrence of rTCA cycle genes with RuBisCo in symbiotic and free-living gammaproteobacteria

Genes for enzymes that are specific to the rTCA pathway, that is the ATP citrate lyase (*aclAB* genes), 2-oxoglutarate:ferredoxin oxidoreductase (OGOR, *korABCD* genes), and a putative fumarate reductase (*tfrAB* genes, homologs of genes encoding a thiol:fumarate reductase from *Methanobacterium thermoautotrophicum* (16)), were assumed to occur in only a few symbiotic Gammaproteobacteria. We discovered, using comparative genomics, that these rTCA cycle enzymes also occur in some Chromatiaceae, including the cultivated sulfur oxidizer *Thioflavicoccus mobilis* and a gammaproteobacterial genome from an environmental metagenome(17), (**Fig. 1**). The type II ATP citrate lyases of tubeworm symbionts and *T. mobilis* were likely acquired via horizontal gene transfer from other bacterial clades (**Suppl. Fig. S1**), (6). These gammaproteobacteria also encode either Form I or II RuBisCO, or both (**Suppl. Note 1**).

**Figure 1:**
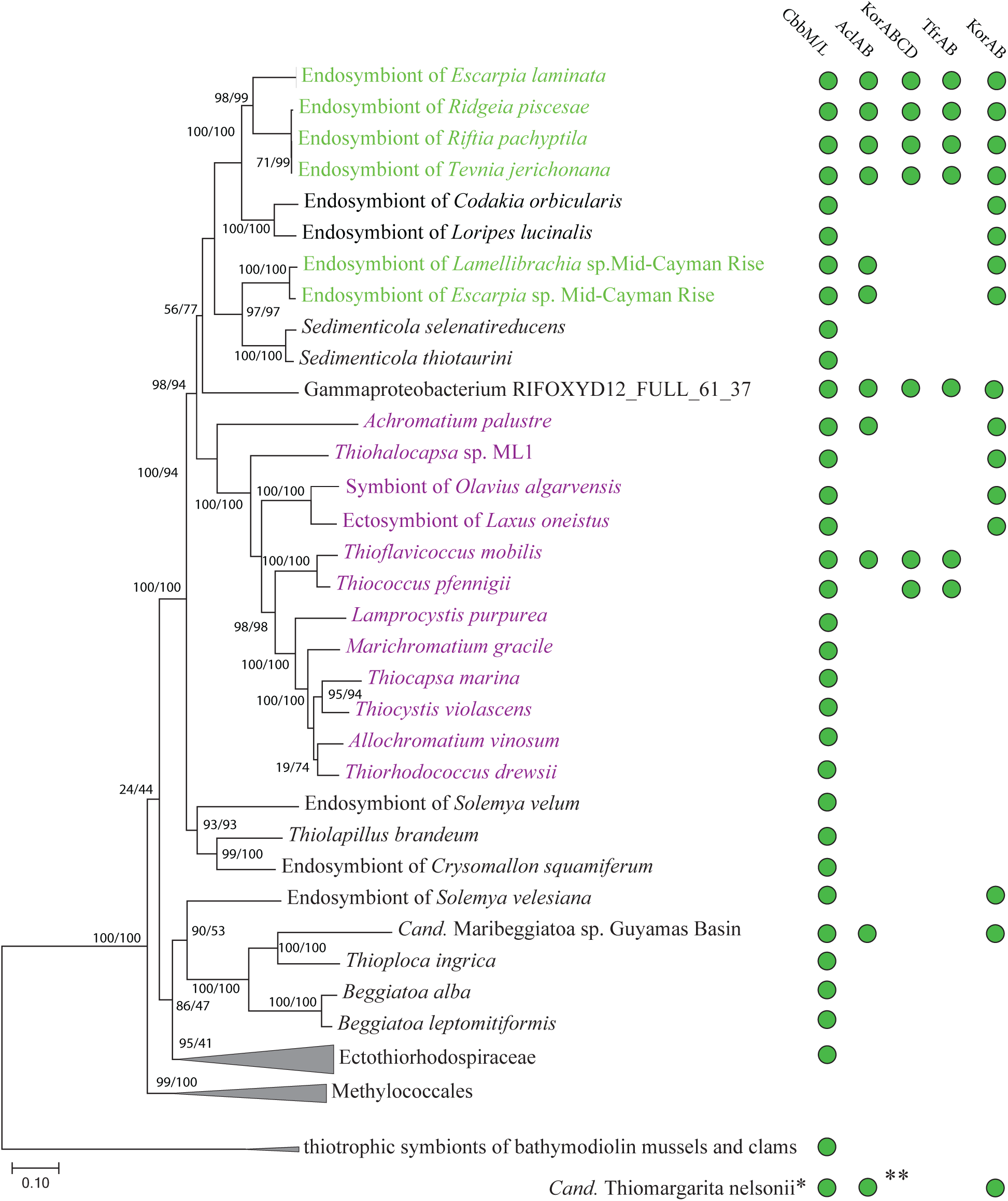
Phylogenomic tree showing occurrence of RuBisCO (CbbM/CbbL), ATP citrate lyase (AclAB), 4-subunit 2-oxoglutarate:ferredoxin oxidoreductase (KorABCD), putative thiol:fumarate reductase (TfrAB) and 2-subunit 2-oxoglutarate:ferredoxin oxidoreductase (KorAB) in the genomes of tubeworm symbionts (green), purple sulfur bacteria (purple) and other related bacteria (58 organisms total, alignment of 2526 amino-acid sites from 23 single-copy markers). The maximum likelihood tree was built with IQ-tree using the LG+R6 model of substitution (20). The tree is unrooted although outgroup “thiotrophic symbionts of bathymodiolin mussels and clams” is drawn at root. Branch labels are SH-aLRT support (%) / ultrafast bootstrap support (%). Accession numbers are provided in Supplementary Table 2. * Was not included in the tree due to several missing single-copy marker genes or multiple versions of these genes, making an accurate phylogenomic placement challenging. ** Only the *aclB* gene was present.

### Presence of the rTCA and the CBB pathways in the genome of a single bacterium

Due to the fragmented nature of the previously available genomes of tubeworm symbionts, past studies could not determine whether the genes for both pathways are present in a single genome or if the two pathways are distributed in a strain-specific manner, i.e. only one of the two pathways is present in the genome of a single cell (3). Here, we provide two lines of evidence that both pathways can co-occur in the genome of a single organism. First, sequencing coverage for the genes of both pathways in the *E. laminata* symbiont was similar to that of single-copy marker genes (**Suppl. Table 1**). Since genes that are strain specific are expected to have lower coverage than the rest of the genome (18), the similar coverage of genes encoding the two pathways and single-copy genes suggests that in the *E. laminata* symbiont both pathways are present in all cells. Second, in the closed genome of the cultured *T. mobilis*, both the genes encoding the rTCA and the CBB cycle co-occur, providing evidence that these genes co-exist in a single genome.

Our transcriptomic analyses of *E. laminata* tubeworm symbionts revealed high expression levels of both the rTCA and the CBB cycle genes (Fig. 2). This observation is consistent with previous proteomic analyses of the *Riftia* symbiont (4, 5). The high expression levels of genes from the rTCA and the CBB cycle suggests that both pathways play an important metabolic role in these symbionts. It is, however, not clear whether these cycles function simultaneously within single symbiont cells, or are differentially expressed within the symbiont population (3).

**Figure 2:**
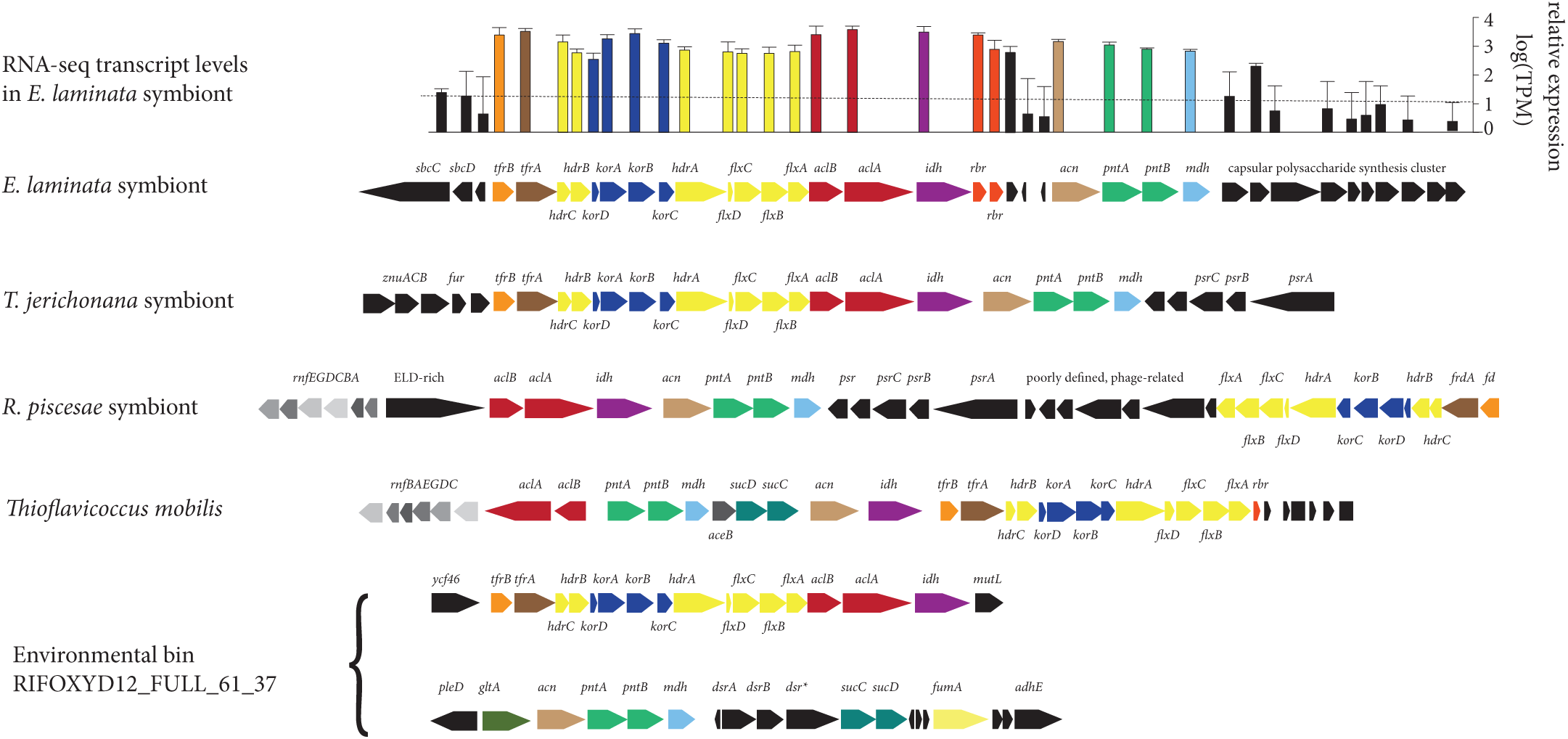
The rTCA cycle gene clusters in symbiotic and free-living bacteria, and the respective transcriptomic gene expression levels in the symbionts of *Escarpia laminata* tubeworm (*aclA*, log (TPM)=3.6; *korA*, log (TPM)=3.3; *hdrA*, log (TPM)=2.9; for comparison - *atpB*, log (TPM)=2.0; *cbbM*, log (TPM)=5.0. TPM, transcripts per kilobase million. *rbr*, rubrerythrin. *dsr\**, oxidoreductase related to the NADPH-dependent glutamate synthase small chain, clustered with sulfite reductase. The dotted line is the median expression value for *E. laminata* genes.

### The rTCA gene clusters are conserved among the tubeworm symbionts and some Chromatiaceae bacteria

In the tubeworm symbionts, the cultivated *T. mobilis* and the gammaproteobacterial genome from an environmental metagenome, there was a considerable level of conservation of the rTCA gene clusters, at the sequence and synteny levels (**Fig. 2**). The *aclAB* genes that encode the two subunits of the ATP citrate lyase were accompanied by those that encode bidirectional TCA cycle enzymes, including *am* (aconitase), *idh* (isocitrate dehydrogenase), and *mdh* (malate dehydrogenase). The other rTCA specific genes *korABCD* (four-subunit OGOR) and *tfrAB* (putative thiobfumarate reductase), were also present in the rTCA gene cluster. Similar to the ATP citrate lyase, the four-subunit OGOR, as well as the thiol: fumarate reductase are very rare among Gammaproteobacteria, and were probably acquired via a single horizontal gene transfer event from a distant bacterial clade (**Fig. 1, Suppl. Fig. S2 and S3**). A dimeric OGOR (*korAB* genes), more common than the four-subunit enzyme among gammaproteobacterial autotrophs, yet absent in *T. mobilis*, was located elsewhere in the genome of the *E. laminata* symbiont. The *korAB* genes were co-localized with genes that encode additional rTCA cycle enzymes (**Suppl. Note 2, Suppl. Fig. S4**).

An array of genes that encode several electron translocating complexes were integrated into the rTCA cycle gene clusters. These complexes included an electron-bifurcating NADH dehydrogenase/heterodisulfide reductase complex (*flxABCD-hdrABC* genes, **Suppl. Note 3**), a NAD(P)+ transhydrogenase and Na+ translocating Rnf membrane complex (*pntAB* and *rnfABCDGE* genes, **Suppl. Note 4**). Most interestingly, the conserved interspersing of the *korABCD* and *tfrAB* genes with the *flxABCD-hdrABC* genes hints at the possibility that these proteins form a complex that efficiently shuttles electrons directly to the OGOR and the thiol: fumarate reductase (**Suppl. Figure S5**). If this is the case, the carbon fixation efficiency of the rTCA cycle would be most likely considerably higher than the canonical rTCA cycle.

## Conclusions

Until now, the only bacteria known to possess two carbon fixation pathways were sulfur-oxidizing, tubeworm symbionts, and possibly also large sulfur bacteria, all of which are currently not amenable to cultivation-based studies. With the discovery of the co-existence of the CBB and rTCA cycle in the cultivable *T. mobilis*, experimental studies are now feasible. Such studies would reveal if these pathways are expressed under different physicochemical conditions, and potentially allow the biotechnological optimization of efficiency and yield in production processes that rely on autotrophic carbon fixers. To our knowledge, the use of organisms with multiple carbon fixation pathways has not been used as a design principle for these applications.

## Methods

### Comparative genomics and transcriptomics

Publically available genomes from NCBI and JGI-IMG collections, as well as de-novo assembled genomes of *Escarpia laminate* symbionts (estimated completeness 99.5%), were used for genomic comparison (see Supplementary Methods). To verify presence/absence of target gene homologs in sequenced organisms we used NCBI’s BLAST against the nucleotide collection and non-redundant protein database (19). *E. laminata* symbiont genomes were used as a template for genome-centered transcriptomics (sequences available under the BioProject accession number PRJNA471406).

### Phylogenetic and phylogenomic analyses

Phylogenomic treeing was performed using scripts available at phylogenomics-tools (DOI:10.5281/zenodo.46122). Twenty-three marker proteins that are universally conserved across the bacterial domain were extracted from genomes using the AMPHORA2 pipeline (20). Twenty-three single-copy markers were used for alignment with MUSCLE (21). The marker alignments were concatenated into a single partitioned alignment, and poorly aligned regions were removed. Functional protein sequences were aligned with MAFFT (22). Maximum Likelihood trees were calculated with IQ-tree (23) and MEGA7 (24), using the best-fitting model.

## Author Contributions

MRB, ND and MK conceived the study. MRB and MK analyzed the samples. MRB, ND and MK wrote the manuscript.

## Acknowledgments

The authors thank all individuals who helped during the R/V Meteor research cruise M114, including onboard technical and scientific personnel, the captain and crew, and the ROV MARUM-Quest team. We thank the Max Planck-Genome-Centre Cologne (http://mpgc.mpipz.mpg.de/home/) for generating the metagenomic and the metatranscriptomic data used in this study. We thank Julie Reveillaud for providing genomes of Mid-Cayman Rise tubeworm symbionts, and Matthias Winkel, Marc Mufimann and Jake V. Bailey for helpful and supportive comments on a bioRxiv preprint version of this manuscript. The Campeche Knoll cruise was funded by the German Research Foundation (DFG – Deutsche Forschungsgemeinschaft). We are grateful to the Mexican authorities for granting permission to conduct this research in the southern Gulf of Mexico (permission of DGOPA: 02540/14 from 5 November 2014). This study was funded by the Max Planck Society, the MARUM DFG-Research Center / Excellence Cluster “The Ocean in the Earth System” at the University of Bremen, an ERC Advanced Grant (BathyBiome, 340535), a Gordon and Betty Moore Foundation Marine Microbial Initiative Investigator Award to ND (Grant GBMF3811) and the NC State Chancellor’s Faculty Excellence Program Cluster on Microbiomes and Complex Microbial Communities (MK).

## Conflict of interest

The authors declare no conflict of interest.

